# MiR-339-3p aggravates rat vascular inflammation induced by AT1R autoantibodies by down-regulating BKα protein expression

**DOI:** 10.1101/2021.10.17.464722

**Authors:** Yang Li, Yan Sun, Mingming Yue, Ming Gao, Li Wang, Ye Wu, Xiaochen Yin, Suli Zhang, Huirong Liu

## Abstract

The abnormality of large-conductance calcium-activated potassium channels (BK channels) is an important factor in inducing vascular inflammation. BK channel agonists can readily recover BK channel function and improve vascular inflammation. However, it is not clear how to improve BK dysfunction caused by downregulation of BK channel protein expression. This study found that angiotensin II-1 receptor autoantibodies (AT1-AA), which are widely present in the body of various types of cardiovascular diseases, can down-regulate the expression of BK channel protein and induce vascular inflammation. Further research found that the elevated neural precursor cells expressed developmentally downregulated 4-like (NEDD4L) protein level is involved in the down-regulation of BK channel α subunit (BKα) protein level by AT1-AA. Bioinformatics analysis and experiments have confirmed that miR-339-3p plays an irreplaceable role in the high expression of NEDD4L and the low expression of BKα, and aggravates the vascular inflammation induced by AT1-AA. Overall, AT1-AA increased miR-339-3p expression (targeting BKα via the miR-339-3p/NEDD4L axis or miR-339-3p alone), reduced BKα protein expression in VSMCs, and induced vascular inflammation. The results of the study indicate that miR-339-3p may become a new target for reversing vascular inflammation in AT1-AA-positive patients.

## Introduction

Vascular inflammation is the pathological basis of various cardiovascular diseases [1]. Inflammatory diseases such as hypertension, atherosclerosis and diabetes are closely related to changes in the expression and function of large-conductance calcium-activated potassium channel (BK channel) [2]. BK channel is the kind of K^+^ channel with the highest expression, the widest distribution, and the largest conductance in vascular smooth muscle cells (VSMCs) [3, [4]. Studies have shown that abnormal BK channel function is involved in the occurrence and development of various inflammations [5, [6]. BK channel agonists can easily restore the function of BK channel and improve vascular inflammation [5, [7]. However, how to improve BK dysfunction caused by the down-regulation of BK channel protein expression has not been understood. Therefore, the factors that reduce the expression of BK channels in VSMCs need to be further explored.

The BK channel is known to be closely related to the renin angiotensin aldosterone system (RAAS) [8, [9]. Overactivation of angiotensin II-1 receptor (AT1R) has proven to be an important reason for the downregulation of BK channel expression in VSMCs [10, [11]. Angiotensin II-1 receptor autoantibody (AT1-AA) is an agonist-like autoantibody that continuously activates AT1R and exerts a vasoconstrictive effect [12]. Studies have shown that AT1-AA is prevalent in vascular inflammation-related diseases, e.g., hypertension [13] and coronary heart disease [14]. However, whether AT1-AA can induce vascular inflammation by decreasing BK channel expression in VSMCs and the underlying molecular mechanisms remains unknown.

Increased protein degradation is an important reason for the decrease in protein expression. Among many pathways of protein degradation, the ubiquitin-proteasome pathway is responsible for 80%-90% of the turnover of intracellular proteins and plays a more important role [15, [16]. Besides, cellular autophagy and apoptosis are also common forms of protein degradation. However, the pathway by which AT1-AA down-regulates the expression of BK channel protein in VSMCs is not clear. This study screened the pathways involved in AT1-AA down-regulating BK channel protein expression in VSMCs and further explored the possible molecular mechanism in the process of AT1-AA down-regulating the expression of BK channel protein.

## Results

### 1. AT1-AA reduced the expression of BKα protein in rat thoracic aortic VSMCs through AT1R

Rats were actively immunized with AT1R-ECII for 12 weeks. The results showed that the OD value of AT1-AA in the serum of AT1R-ECII group rats was significantly higher than the OD value of AT1-AA in the serum of saline group rats (Suppl. Figure 1A), suggesting that the AT1-AA active immunization rat model was successfully established. Further, the systolic and diastolic arterial blood pressures of AT1-AA-positive rats were found to be significantly increased (Suppl. Figure 1B), and ultrasound results showed a significant increase in the thickness of the thoracic aortic vessel wall (Suppl. Figure 1C), indicating that the blood vessels have a damaged phenotype. To verify the effect of AT1-AA on the expression of BK channel proteins in VSMCs, we detected the expression of BK protein by Western blot and immunofluorescence. The results showed that compared with the saline group, the BKα protein level in the thoracic aorta of AT1-AA-positive rats was significantly reduced (Figure. 1A-B), but there was no significant difference in the BKβ1 protein level compared with the saline group (Suppl. Figure 1G-H). VSMCs treated with AT1-AA also showed similar results (Figure. 1C and Suppl. Figure 1I).

**Figure 1.**
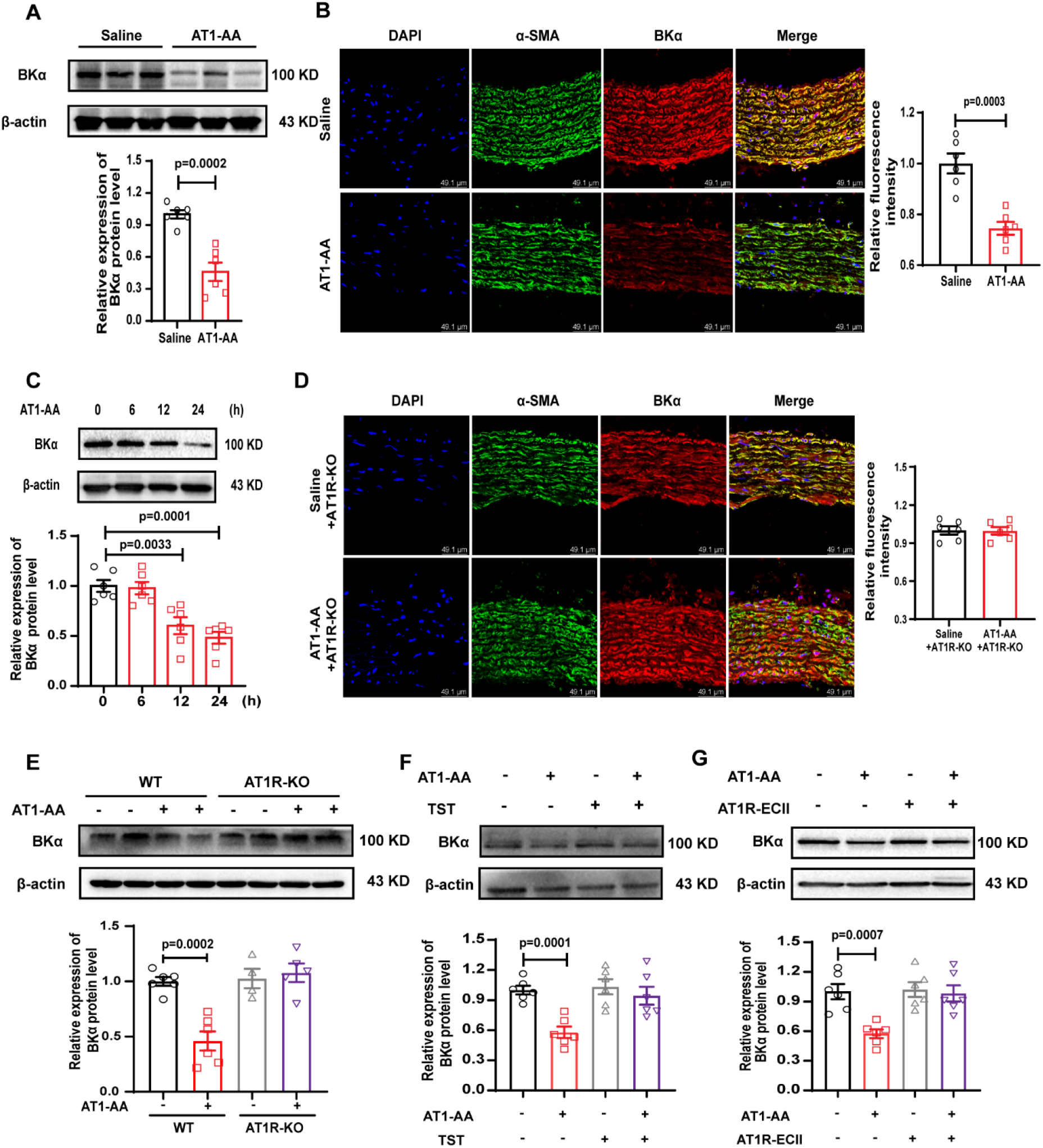
AT1-AA affected the expression of BKα protein in rat thoracic aortae and VSMCs. Western blot (A) and immunofluorescence (B) were used to detect the BKα protein level in the thoracic aorta of AT1-AA-positive rats, bar=49.1 μm, n=6. (C) Changes in BKα protein levels in VSMCs caused by AT1-AA at different times were detected, n=6. Immunofluorescence (D) and Western blotting (E) were used to detect the changes in BKα protein levels in the thoracic aorta of AT1-AA-positive AT1R knockout rats, bar=49.1 μm, n=4, 6. (F) After pretreating VSMCs with telmisartan to block AT1R, the changes in BKα protein levels caused by AT1-AA were detected (n=6). (G) Western blotting was used to detect the BKα protein level after treating the VSMCs with the mixture for 24 h, in which AT1R-ECII was premixed with AT1-AA, n=6. The results of each sample were tested three times.

To further verify whether the decreased BKα protein expression in VSMCs induced by AT1-AA was dependent on the AT1R pathway, we found that AT1-AA did not affect the protein level of BKα in the thoracic aorta of AT1R-knockout rats following active immunization (Figure. 1D-E). In addition, there was no significant change in BKα protein expression when VSMCs were pretreated with the AT1R blocker telmisartan (TST) or antigen peptide AT1R-ECII (Figure. 1F-G). The above results indicated that AT1-AA downregulated the protein expression of BKα in VSMCs via an AT1R-dependent pathway.

### 2. Downregulation of BKα protein in VSMCs promoted the vascular inflammatory response induced by AT1-AA

CD3, CD19 and CD68 were used to label T and B lymphocytes and macrophages, respectively. Immunofluorescence staining revealed that inflammatory cells evidently increased in the middle layer of the thoracic aorta of AT1-AA-positive rats (Figure. 2A and Suppl. Figure 2A). Meanwhile, the protein expression of inflammatory cytokines (including IL-6, IL-1β and TNF-α) in the thoracic aortas of AT1-AA-positive rats also increased significantly (Figure. 2B). After treatment of primary cultured rat thoracic aortic VSMCs with AT1-AA, the protein expression of inflammatory cytokines in cells (Figure. 2C) and cell supernatant (Figure. 2D) apparently increased. Correlation analysis found that the decreased BKα protein level induced by AT1-AA was significantly related to the high expression of inflammatory cytokines in VSMCs (Figure. 2E).

**Figure 2.**
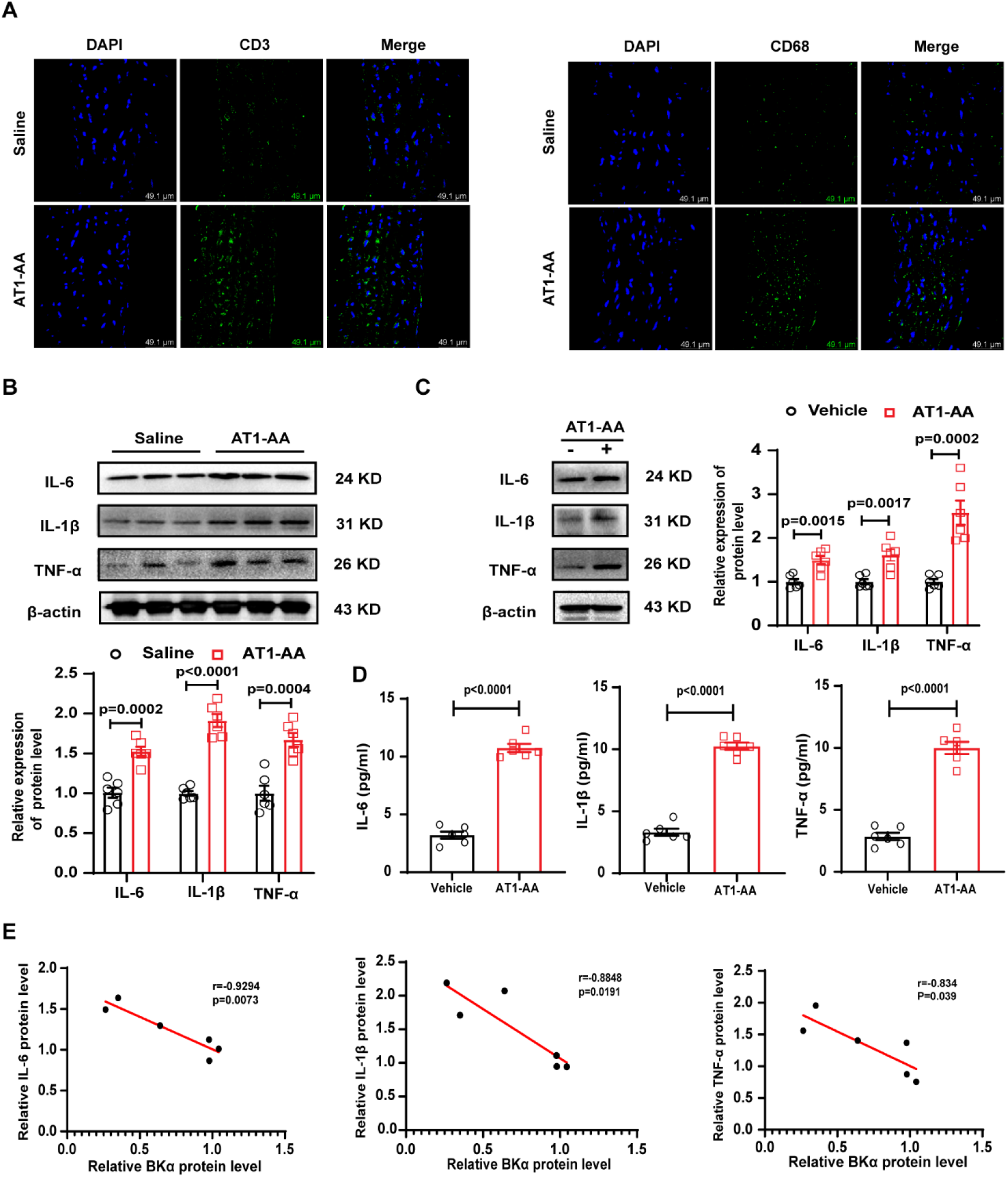
The high expression of inflammatory cytokines induced by AT1-AA is negatively correlated with the level of BKα protein. (A) Immunofluorescence was used to detect the expression of CD3 and CD68 in the thoracic aortic vessel wall of AT1-AA-positive rats, bar=49.1 μm. The protein expression of inflammatory cytokines (including IL-6, IL-1β and TNF-α) in (B) the thoracic aorta of AT1-AA-positive rats, and (C) VSMCs treated with AT1-AA were detected, n=6. (D) To detect the protein expression of inflammatory cytokines (including IL-6, IL-1β and TNF-α) in the culture supernatant of VSMC after AT1-AA treatment though ELISA, n= 6. (E) The relationship between the downregulation of BKα protein levels and the high expression of inflammatory cytokines induced by AT1-AA in vitro was analysed. The results of each sample were tested three times.

To prove that low expression of BKα protein in VSMCs was involved in AT1-AA-induced vascular inflammation, immunofluorescence results were obtained for the thoracic aorta of BKα-knockout rats, and the results indicated that compared with wild-type rats, the infiltration of inflammatory cells increased significantly in the vascular wall of the thoracic aortas of BKα-knockout rats (Figure. 3A and Suppl. Figure 2F). Moreover, compared with the vehicle group, the IL-6, IL-1β and TNF-α protein levels increased significantly after BKα knockdown in primary VSMCs (Figure. 3B) and cell supernatant (Figure. 3C). The BKα overexpression adenovirus was used to upregulate the BKα protein level in the blood vessels of AT1-AA-positive rats, and the results showed that BKα overexpression was able to reverse the AT1-AA-induced inflammatory cell infiltration in the middle layer of thoracic aorta blood vessels (Figure. 3D and Suppl. Figure 2K) and the expression of inflammatory cytokines in blood vessels (Figure. 3E).

**Figure 3.**
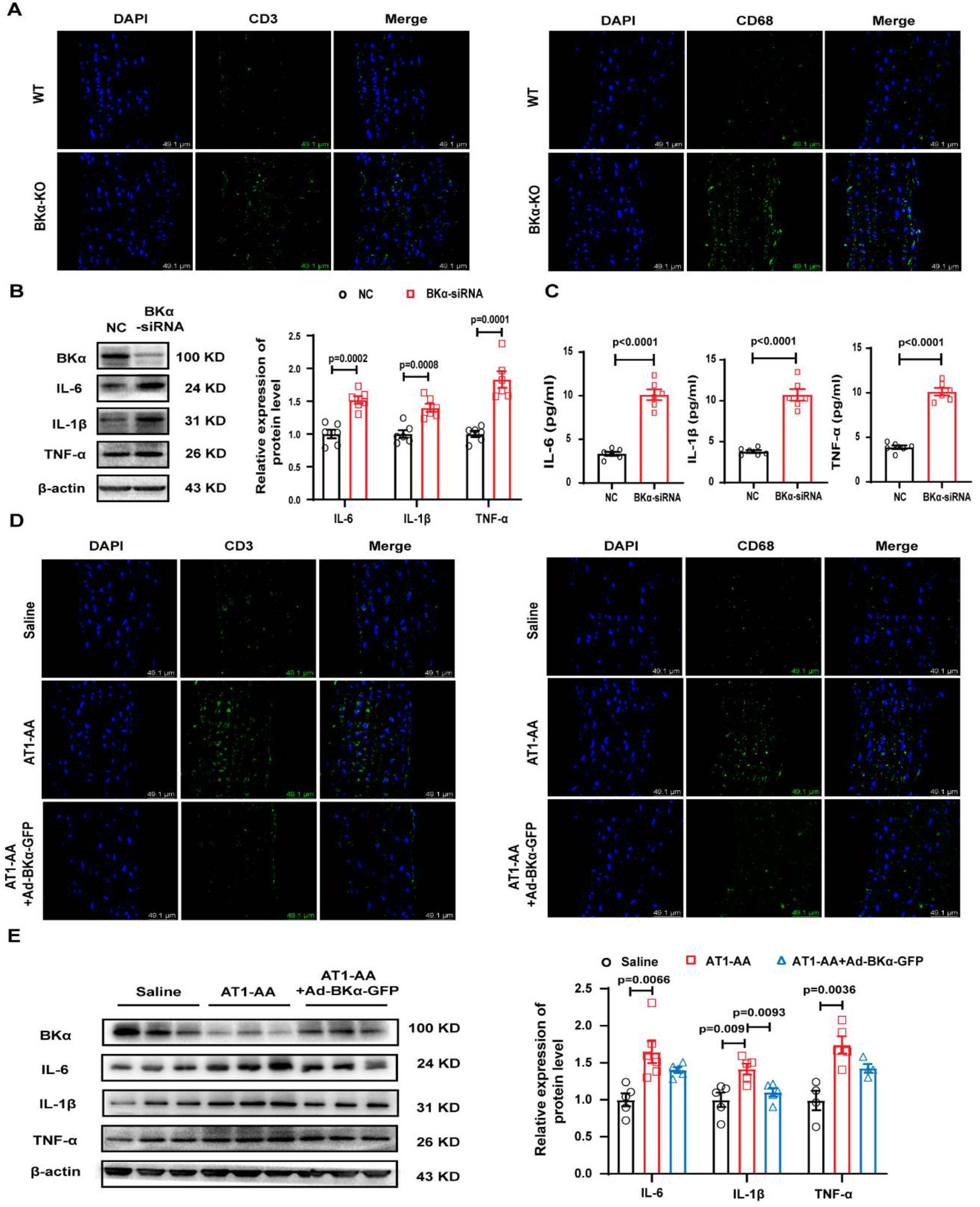
The downregulation of BKα protein levels in VSMCs participated in AT1-AA-induced vascular inflammation. (A) Immunofluorescence was used to observe the expression changes of CD3 and CD68 in the thoracic aortic wall of BKα knockout rats, bar=49.1 μm. Western blot and ELISA were used to detect the protein expression of IL-6, IL-1β and TNF-α in (B) primary VSMCs, and (C) cell supernatant after knocking down BKα, n=6. (D) After overexpression of BKα, immunofluorescence was used to observe the changes in inflammatory cell marker molecules (bar=49.1 μm). (E) Western blot was used to detect the reverse effect of BKα overexpression on the increased expression of inflammatory cytokines induced by AT1-AA in vivo, n=4, 5. The results of each sample were tested three times.

### 3. AT1-AA could reduce the expression of BKα by increasing the ubiquitin-related protein NEDD4L in VSMCs

Using RT-PCR to detect the effect of AT1-AA on the BKα mRNA level of AT1-AA-positive rat aortas and VSMCs treated with AT1-AA, we found that AT1-AA did not change the BKα mRNA level (Suppl. Figure 3A and Figure. 4A), suggesting that AT1-AA cannot affect the BKα transcription of VSMCs. The posttranslational modification of proteins is an important regulatory mechanism that affects the protein level [17]. Therefore, the ubiquitin proteasome pathway inhibitor MG-132, autophagy inhibitor 3-MA, and apoptosis inhibitor Z-VAD-FMK were used to verify which pathway was involved in the reduction in BKα protein expression, and MG-132 obviously reversed the downregulation of BKα protein expression in VSMCs induced by AT1-AA (Figure. 4B and Suppl. Figure 3B), demonstrating that the ubiquitin pathway was largely involved in the decrease in BKα protein levels induced by AT1-AA. Subsequently, the CoIP method was used to confirm that AT1-AA can directly induce a significant increase in the ubiquitination level of BKα in VSMCs (Figure. 4C). Next, the E3 ligase NEDD4L was found to possibly be the ubiquitin-related protease involved in the decrease in BKα protein levels by AT1-AA through protein profile analysis (Suppl. Figure 3C-D). AT1-AA markedly increased the NEDD4L protein level in VSMCs, as shown by Western blot analysis (Figure. 4D). The immunofluorescence results were consistent with the Western blot results; compared with the vehicle group, and the red fluorescence-labelled NEDD4L was significantly increased in the VSMCs treated with AT1-AA (Figure. 4E). We identified that AT1-AA can significantly increase the protein level of NEDD4L bound to BKα in VSMCs using the CoIP method, and it was further confirmed that under AT1-AA treatment, the interaction between NEDD4L and BKα evidently increased (Figure. 4F). After knocking down NEDD4L in VSMCs and then giving AT1-AA treatment, the phenomenon of AT1-AA decreasing the expression of BKα was found to disappear (Figure. 4G). The above results suggested that AT1-AA promoted the interaction between NEDD4L and BKα protein by increasing the protein level of NEDD4L, thereby downregulating the expression of BKα protein in VSMCs.

**Figure 4.**
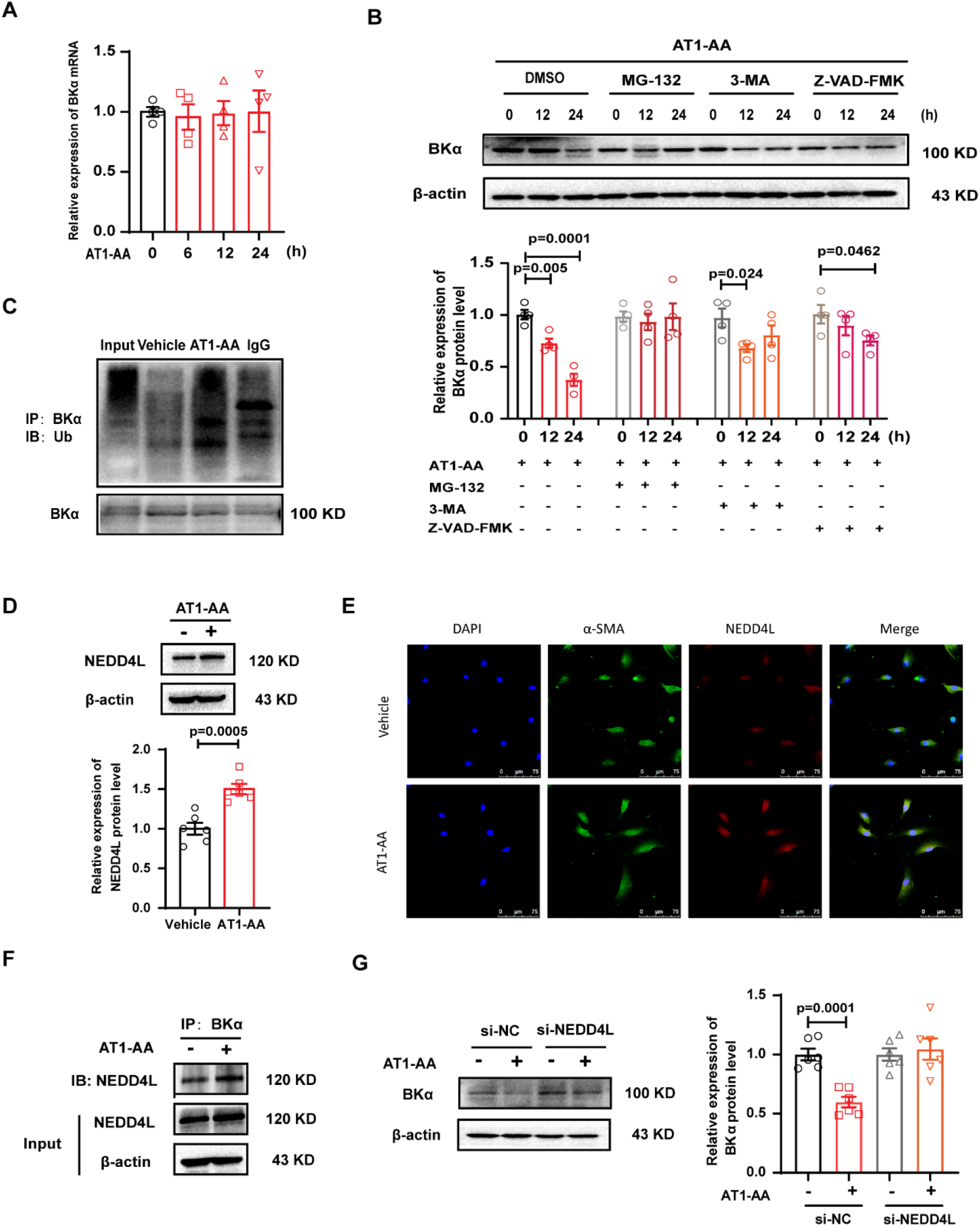
NEDD4L participated in AT1-AA downregulating BKα protein expression in VSMCs. (A) RT-PCR was used to detect BKα mRNA levels in VSMCs after AT1-AA treatment, n=4. (B) The possible pathway which AT1-AA downregulates the BKα protein level in VSMCs was detected by Western blot, n=4. (C) CoIP was used to detect the effect of AT1-AA on the level of BKα ubiquitination. The effect of AT1-AA on NEDD4L protein levels in VSMCs was proven by (D) Western blot and (E) immunofluorescence, bar=75 μm, n=6. (F) The protein level of NEDD4L connected with BKα after AT1-AA treatment to VSMCs was detected by the CoIP. (G) Western blot was used to observe the effect of AT1-AA on BKα protein level after knocking down NEDD4L, n=6. The results of each sample were tested three times.

### 4. MiR-339-3p inhibited the expression of BKα in VSMCs by upregulating NEDD4L

Numerous studies have shown that if the microRNA combines with the 5’UTR of the target gene mRNA, the expression of the target gene can be promoted [18], and if the microRNA combines with the 3’UTR of the target gene mRNA, it will cause the degradation of the mRNA or inhibit the translation of the target mRNA. To explore the mechanism by which AT1-AA increased NEDD4L protein expression and downregulated BKα protein expression in VSMCs, three miRNAs were screened out that simultaneously target the 5’UTR of NEDD4L and the 3’UTR of BKα through bioinformatics analysis (http://mirwalk.umm.uni-heidelberg.de/), including miR-145-5p, miR-149-5p and miR-339-3p (Figure. 5A). The expression of miR-339-3p was also found to increase most significantly after AT1-AA treatment of VSMCs (Figure. 5B). The expression of miR-339-3p in the thoracic aortas of AT1-AA-positive rats also increased significantly (Figure. 5C). Meanwhile, fluorescence in situ hybridization also showed that the expression of miR-339-3p in the cytoplasm of VSMCs treated with AT1-AA was significantly higher than the expression of miR-339-3p in the cytoplasm of the vehicle group (Figure. 5D). The above results indicated that AT1-AA can increase the expression of miR-339-3p in VSMCs. To further prove that miR-339-3p can affect the protein expression of NEDD4L and BKα in VSMCs, we used bioinformatics analysis to identify potential matching sites between miR-339-3p and the two proteins and found that there were binding sites for rat miR-339-3p in the 5’UTR of NEDD4L (Figure. 5E) and the 3’UTR of BKα (Figure. 5I).

**Figure 5.**
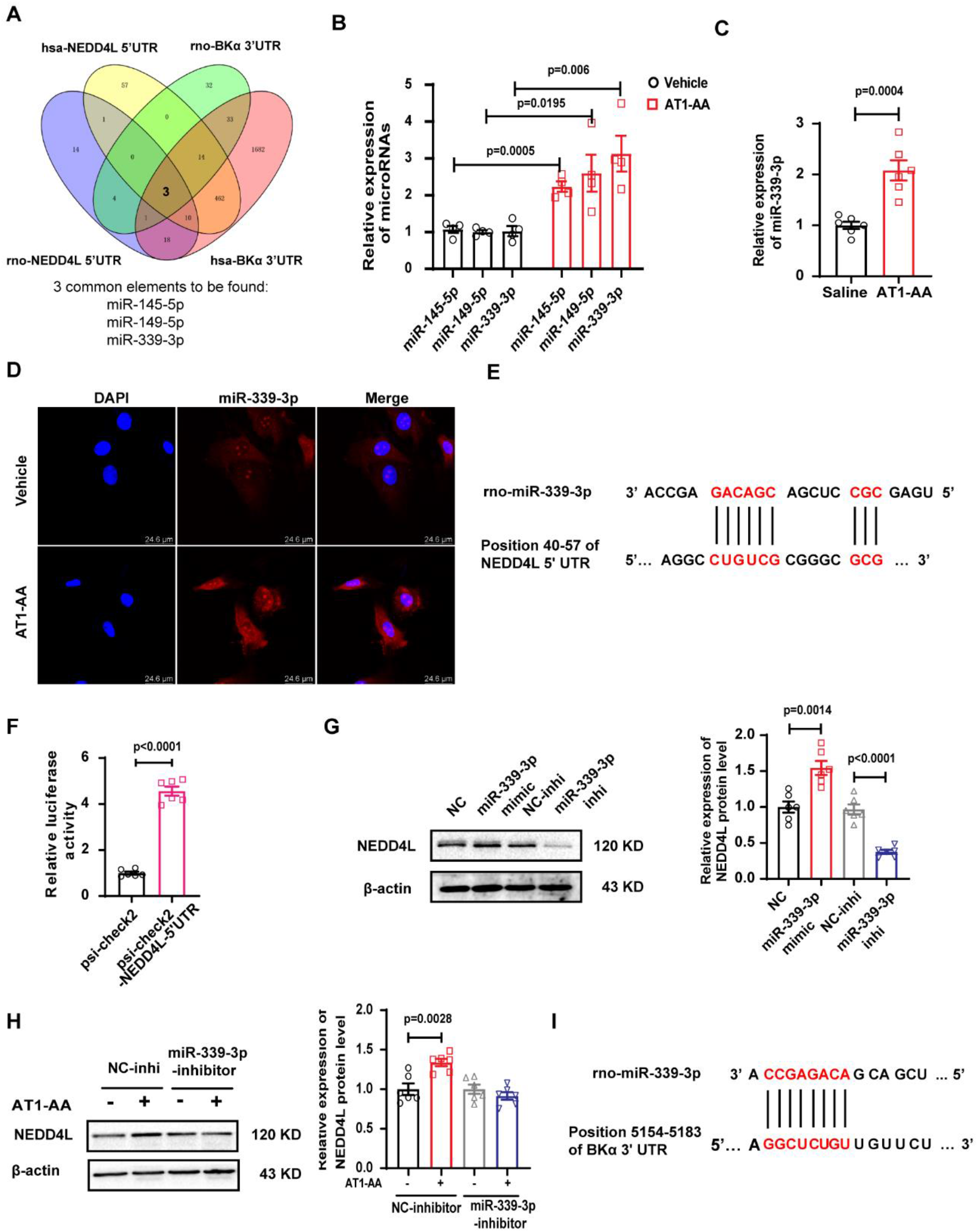
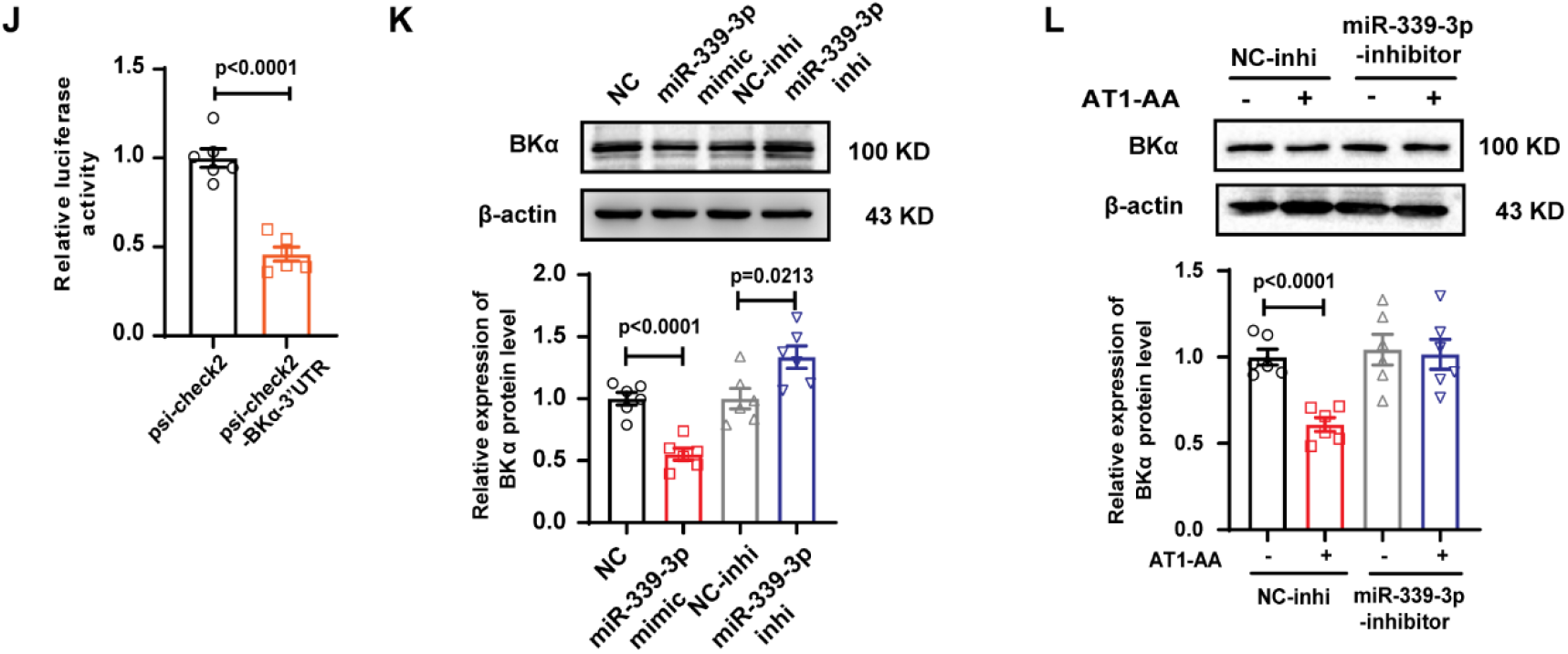
miR-339-3p reduced the expression of BKα in VSMCs by upregulating NEDD4L. (A) miRNAs targeting the NEDD4L 5’UTR and BKα 3’UTR were analysed by bioinformatics (http://mirwalk.umm.uni-heidelberg.de/). (B) The expression of different miRNAs in AT1-AA-treated VSMCs and (C) the expression of miR-339-3p in the thoracic aorta of AT1-AA-positive ratswere detected by RT-PCR, n=4. 6. (D) FISH was used to observe the level of miR-339-3p in VSMCs treated with AT1-AA; bar=24.6 μm. (E) The binding sites of miR-339-3p and the NEDD4L 5’UTR were analysed by bioinformatics. (F) The effective binding of miR-339-3p and the NEDD4L 5’UTR was proven by the luciferase reporter gene method, n=6. Western blot was used to detect (G) the changes in NEDD4L protein levels after overexpression/inhibition of miR-339-3p, and (H) after inhibiting miR-339-3p, the NEDD4L protein level was detected in VSMCs treated with AT1-AA (n=6). The binding sites of miR-339-3p and the BKα 3’UTR were analysed by bioinformatics (I). (J) The effective binding of miR-339-3p and the BKα 3’UTR was proven by the luciferase reporter gene method, n=6. (K) Western blot was used to detect the changes in BKα protein levels after overexpression/inhibition of miR-339-3p, and (L) the inhibition of miR-339-3p before treatment with AT1-AA was used to detect BKα protein levels, n=6. The results of each sample were tested three times.

To verify the effective combination of miR-339-3p and the NEDD4L 5’UTR and evaluate the effect of miR-339-3p on NEDD4L expression, we constructed a psi-check 2-NEDD4L-5’UTR plasmid (Suppl. Figure 4B), and it was cotransfected with miR-339-3p mimic into HEK293A cells. The results of the dual luciferase reporter assay showed that compared with psi-check 2, miR-339-3p mimics and psi-check 2-NEDD4L-5’UTR cotransfection significantly increased the luciferase activity (Figure. 5F). Then, we overexpressed or knocked down miR-339-3p in VSMCs and observed the protein expression of NEDD4L. The results showed that overexpression of miR-339-3p significantly increased the expression of NEDD4L protein, while knockdown of miR-339-3p significantly decreased the expression of NEDD4L protein (Figure. 5G). When VSMCs were treated with AT1-AA after knockdown of miR-339-3p, the protein level of NEDD4L evidently did not change (Figure. 5H), suggesting that miR-339-3p was an important mechanism of AT1-AA-induced changes in NEDD4L protein levels.

To verify the effective binding of miR-339-3p to the BKα 3’UTR and evaluate whether miR-339-3p has a direct effect on the expression of BKα, we also constructed a psi-check 2-BKα-3’UTR plasmid (Suppl. Figure 4C) and cotransfected the plasmid with miR-339-3p mimic into HEK293A cells. The results showed that miR-339-3p mimic and psi-check 2-BKα-3’UTR cotransfection significantly decreased luciferase activity (Figure. 5J). Overexpression of miR-339-3p was also found to decrease the level of BKα protein, while inhibiting miR-339-3p induced an increase in BKα protein levels (Figure. 5K). After miR-339-3p was inhibited and then treated with AT1-AA, the decrease of BKα protein levels in VSMCs induced by AT1-AA disappeared (Figure. 5L), suggesting that miR-339-3p was also one of the reasons for the downregulation of BKα protein expression in VSMCs induced by AT1-AA.

The above results suggested that the increase in miR-339-3p expression in VSMCs induced by AT1-AA not only promoted the decrease in BKα expression by upregulating the NEDD4L protein level but also directly decreased the protein expression of BKα in VSMCs.

### 5. Inhibition of miR-339-3p can reverse vascular inflammation induced by AT1-AA in vivo

We tried to use antagomir-339-3p to reverse the decrease in BKα protein levels and vascular inflammation in AT1-AA-positive rats. First, to prove the effectiveness of antagomir-339-3p, VSMCs were transfected with antagomir-339-3p, and miR-339-3p expression in VSMCs was significantly reduced (Suppl. Figure 5A). Similarly, antagomir-339-3p successfully inhibited the expression of miR-339-3p in SD rats through tail vein injection (Figure. 6B). Then, the BKα protein level and the protein expression of inflammatory cytokines in rat thoracic aortas were detected. Compared with the AT1-AA group, treatment with antagomir-339-3p reversed the decreased BKα protein level (Figure. 6C-D) and lessened the infiltration of inflammatory cells in the rat thoracic aortas induced by AT1-AA via immunofluorescence (Figure. 6E). Finally, we found that compared with the AT1-AA group, antagomir-339-3p injection reversed the increased expression of the inflammatory cytokines IL-6, IL-1β and TNF-α in the rat thoracic aortas (Figure. 6F). The results of small animal ultrasound showed that after the AT1-AA-positive rats were injected with antagomir-339-3p, the thickness of the thoracic aortic wall of the rats was significantly improved compared with the rats in the AT1-AA group (Suppl. Figure 5B). In addition, from recorded arterial blood pressure data of these model rats, we found that antagomir-339-3p can alleviate the increase in systolic and diastolic arterial blood pressure caused by AT1-AA (Suppl. Figure 5C). The above results suggested that inhibiting miR-339-3p can significantly reverse the vascular inflammation induced by AT1-AA.

**Figure 6.**
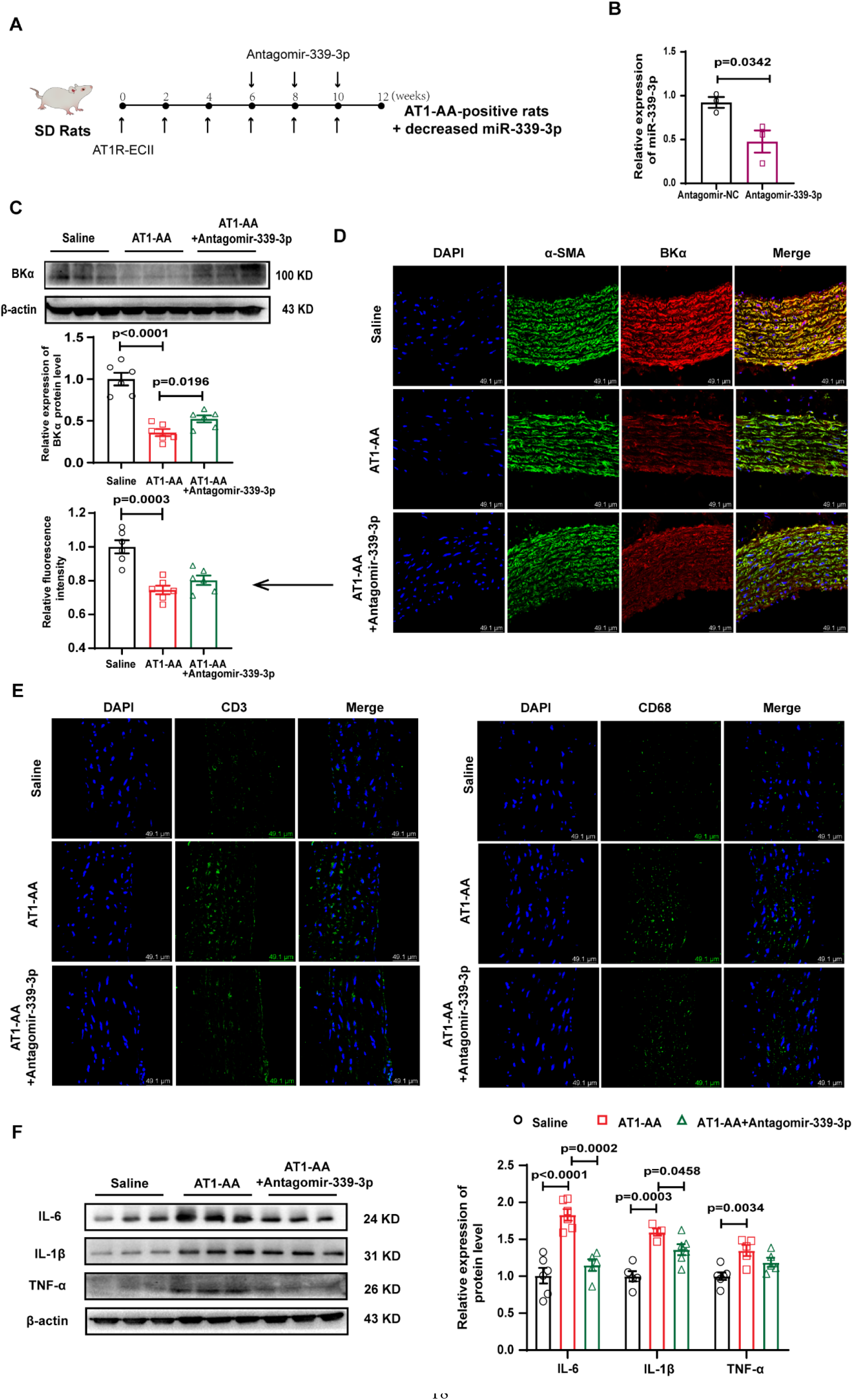
Changes in AT1-AA-induced vascular inflammation after inhibiting miR-339-3p. (A) Model animal production process. (B) The level of miR-339-3p in the thoracic aorta of SD rats injected with antagomir-339-3p was detected by RT-PCR, n=3. (C) Western blot and (D) immunofluorescence were used to detect changes in BKα protein levels in the thoracic aorta vascular wall of rats after injection of antagomir-339-3p into AT1-AA-positive rats, n=6. (E) Immunofluorescence was used to observe the surface marker molecules of AT1-AA-positive rat thoracic aortic inflammatory cells injected with antagomir-339-3p; bar=49 μm. (F) Detection of the expression of the inflammatory cytokines IL-6, IL-1β and TNF-α in the vascular wall of the rat thoracic aorta after the injection of antagomir-339-3p in AT1-AA-positive rats by Western blot, n=5-6. The results of each sample were tested three times.

## Discussion

In general, VSMCs play an important role in the inflammatory process of blood vessel walls. The role and mechanism of VSMCs in the inflammatory process of the aortic wall have not been fully elucidated. Once the body has an inflammatory response, blood vessels serve as conduits through various tissues and organs, which can cause more extensive pathological changes when inflamed, and deliver the produced inflammatory factors to various parts of the body [19]. In this study, we investigated the mechanism of AT1-AA-induced inflammation in the vascular wall, mainly in VSMCs, and attempt to identify new targets that may reverse inflammation in the vascular wall of AT1-AA-positive patients.

Excessive activation of AT1R on the surface of VSMCs has been found to be an important mechanism of vascular inflammation [20]. Using various drugs of the sartan family to inhibit AT1R can significantly reduce the occurrence of atherosclerosis in mice, rabbits and monkeys [21]. As a persistent agonist of AT1R, AT1-AA exists widely in the serum of patients with cardiovascular diseases such as hypertension and can cause excessive and continuous activation of AT1R by binding to the extracellular second loop of the AT1 receptor (AT1R-ECII), which can directly damage endothelial cells and VSMCs, leading to an increase in transcription factors related to proinflammatory responses [22]. Once these transcription factors reach the vascular system, they drive and accelerate vascular inflammation [23, [24]. CD3, CD19 and CD68 are the surface markers of T cells, B cells and macrophages, respectively. However, during our experiment, it was found that CD3 and CD68 increased significantly in the blood vessel wall. Compared with this, the increase in CD19 expression was milder. Studies have shown that cellular immunity plays an important role in the pathogenesis of aortitis, and a large number of immune cells such as T cells, macrophages and natural killer cells are mainly infiltrated in the wall specimens of aortic arteritis [25]. IL-1β is a typical proinflammatory cytokine that induces the production of a broad spectrum of cytokines and chemokines, leading to the recruitment of various types of inflammatory cells [26]. The high expression of IL-1β precursor protein can indicate the initiation of inflammatory response [27]. Interleukin-6 (IL-6) is induced by IL-1 and plays a central role in the process of inflammation, which is more sensitive and lasts longer than other cytokines. TNF-α is an effective proinflammatory cytokine that can regulate the expression of various proteins, such as IL-1 and IL-6 [28]. Increased expression of inflammatory cytokines in VSMCs and cell supernatants suggested that AT1-AA can cause inflammatory changes in VSMCs and aggravate vascular inflammation, but the specific mechanism is not fully understood.

Potassium channel disorders play a pivotal role in various diseases with significant inflammatory changes, such as systemic hypertension, diabetes, and atherosclerosis [29]. Potassium channels in smooth muscle cells (SMCs) have been reported to participate in the release of proinflammatory factors by SMCs and play an important role in the inflammatory pathological process of atherosclerosis [30]. In recent years, an increasing number of potassium channels have become potential targets for the treatment of inflammatory diseases [31]. Calcium-activated potassium channels (KCas) are divided into large conductance (BKCa), intermediate conductance (IKCa) and small conductance (SKCa) types. Among these channels, BKCa is expressed mainly in VSMCs [32]. BKCa channels (BK channels) are composed mainly of α subunits that form pores and auxiliary β subunits that regulate channel Ca^2+^ sensitivity, activity and structure, and they play an important role in regulating physiological processes, including smooth muscle tension and neuronal excitability [33]. Studies have shown that blocking BK channel function can promote the occurrence and development of vascular inflammation, and the vascular inflammation caused by ischaemia-reperfusion is partially reversed after activation of the BK channel induced by NS1619 [5]. Our group previously found that AT1-AA can downregulate the function of BK channels and damage blood vessels [34]. In this experiment, we pretreated VSMCs with the BK channel agonist NS1619 to upregulate the function of the BK channel and then treated them with AT1-AA. We found that NS1619 indeed upregulated the activity of the BK channel (Suppl. Figure 2B) and partially reversed the vascular inflammation induced by AT1-AA (Suppl. Figure 2C) but could not achieve complete reversal, suggesting that in addition to the impairment of BK channel function, there are other factors that play an irreplaceable role in AT1-AA-induced vascular inflammation. Protein expression is a necessary condition for its function [35], and studies have shown that overactivation of AT1R leads to a decrease in the expression of BK channels [10], but it is not clear whether AT1-AA induces vascular inflammation by reducing the expression of BK channels. In this experiment, to clarify the specificity of AT1-AA decreasing BKα, we established a negative IgG group, an unrelated antibody group, namely, the β1 adrenergic receptor autoantibody (β1-AA) group, and a positive group (Ang II), and found that only AT1-AA can lower BKα protein levels (Suppl. Figure 2D).

To confirm the role of BKα in AT1-AA-induced vascular inflammation, we observed a significant increase in inflammatory cell infiltration in the thoracic aorta of BKα-knockout rats. However, we observed that the changes in systolic and diastolic arterial blood pressure in 5-month-old BKα-knockout rats were not obvious (Suppl. Figure 2E). Although vessels without BK channels may increase contractility due to calcium flow in VSMCs, we also found that the heart function of BKα-knockout rats was damaged, leading to cardiac contractility and ejection dysfunction. We speculate that this may be the reason why there was no significant change in arterial blood pressure in BKα-knockout rats. However, after BKα overexpressing adenovirus was injected into the tail vein of AT1-AA-positive rats, it was detected that the thoracic aortic wall thickness and arterial blood pressure of AT1-AA-positive rats had a reversal effect (Suppl. Figure 2L-M). Due to the technical difficulties in constructing conditional knockout rats, we used BKα global gene knockout rats in our experiment. With the development of technology, we constructed VSMC-specific knockout BKα rats and put them into followup experiments to prove that the loss of BKα in SMCs is an important cause of vascular inflammation.

It is very important to further explore the molecular mechanism of AT1-AA down-regulating the expression of BKα protein. Our experiments proved that AT1-AA does not affect the transcription level of BKα protein. In addition to transcription, posttranscriptional regulation and posttranslational modification may also be involved in the regulation of protein levels, in which posttranslational modification is an important factor affecting protein expression. Ubiquitin, as the most common type of posttranslational modification, is also an efficient and extensive pathway for protein degradation [36]. Some ubiquitin related enzymes involved in BK channel ubiquitination have been reported, including F-box protein (FBXO) [37], with-no-lysinekinase-4 (WNK4) [38], muscle RING finger protein 1 (MuRF1) [39], and CRL4A (CRBN). Autophagy is also an important intracellular degradation system in which intracellular substances are transported to lysosomes and degraded in lysosomes, dynamically circulating intracellular energy and substances [40]. Apart from this, some proteins can also be degraded during apoptosis, including DNA damage repair enzymes, U1 small nuclear ribonucleoprotein components and actin, etc. With the experimental data, we observed that all three pathways are involved in AT1-AA-induced reduction of BKα channel expression in VSMCs, but in contrast, the ubiquitination pathway plays a more important role in this process (Figure. 4B and Suppl. Figure 3B), and therefore the degradation of the BK channel ubiquitination pathway was the focus of the present study.

The ubiquitin process is a three-enzyme cascade catalytic process that consists of E1-ubiquitin activating enzyme, E2-ubiquitin-binding enzyme and E3-ubiquitin ligase. The interaction between the E3-ubiquitin ligase and target protein is the core step of ubiquitin-mediated protein degradation [41]. In the whole process, the main function of ubiquitin is to mark proteins that need to be decomposed so that they can be hydrolysed. In addition to labelling proteins present in the cytoplasm, ubiquitin can also label transmembrane proteins and remove them from the cell membrane [42]. In this experiment, we screened and identified different ubiquitin-related proteases directly connected to BKα through protein profiling methods on the vehicle group and the VSMCs treated with AT1-AA. The data were analysed by GO, and 32 proteins were found in the “protein binding” category after classification according to molecular function. The biological effects of these proteins were queried by the UniProt database (https://www.UniProt.org/), and only one of them was related to protein degradation (Suppl. Figure 3A). As a result, an E3-ubiquitin ligase named neural progenitor cells expressing developmental downregulated 4-like proteins (NEDD4L or Nedd4-2) was screened out. The main targets of NEDD4L are membrane proteins, including ion channels and transporters [43]. NEDD4L has been reported to be able to negatively regulate the cell surface level of several ion channels, receptors and transporters involved in regulating neuronal excitability [44], targeting mainly voltage-gated sodium channels. Studies have shown that the WW functional domain of NEDD4L interacts with the PY functional domains of various subunits of ENaC, leading to the ubiquitination of ENaC and the endocytosis and degradation of ENaC from the cell membrane [45, [46]. Besides, knockout of NEDD4L can lead to high expression of epithelial Na^+^ channels and eventually aggravate pulmonary inflammation [47], suggesting that NEDD4L is related to inflammation. So how does AT1-AA increase the protein level of NEDD4L?

The increase in mRNA translation is a direct link to the increase in protein levels, and mature microRNAs (miRNAs) can regulate the translation of target mRNA. There are increasing reports about the involvement of miRNAs in the regulation of the ubiquitin-proteasome system (UPS) [48, [49]. In most cases, miRNAs guide the silencing complex (RISC) to degrade mRNA or hinder its translation by pairing with the 3’UTR of the target gene mRNA base [50]. At the same time, some studies have shown that miRNAs can also bind to the 5’UTR of the target gene mRNA and promote the expression of the target gene [18]. In this study, we obtained the miRNA intersection of targeting human and rat NEDD4L 5’UTR and BKα 3’UTR by bioinformatics and selected miR-339-3p as the target miRNA for follow-up verification. According to existing research, miR-339-3p is closely related to the proliferation, migration and invasion of all kinds of cancer cells [51], but its regulatory effect on the UPS has not been reported. This study conclusively confirmed that miR-339-3p targets both the 5′UTR of NEDD4L as well as the 3′UTR of BKα, both achieving reduced BKα protein expression and exacerbating vascular inflammation. We innovatively demonstrate that a single miRNA can simultaneously target different regions of two proteins and lead to subsequent cascade amplification.

## Conclusion

This study demonstrated that the reduction in BKα protein was the key mechanism of vascular inflammation induced by AT1-AA, and the high expression of the E3-ubiquitin ligase NEDD4L was involved in the downregulation of BKα by AT1-AA. MiR-339-3p played an irreplaceable role in both high expression of NEDD4L and low expression of BKα, aggravating the vascular inflammation induced by AT1-AA. From the point of view of ubiquitin degradation and miRNA regulation, this study searched for the molecular mechanism of the AT1-AA-induced downregulation of BKα protein expression in VSMCs and tried to provide a new possible treatment for vascular inflammation-related diseases aggravated by the vascular smooth muscle cell inflammatory phenotype in AT1-AA-positive patients.

## Materials and methods

### Establishment of model animals

Two-hundred-gram 8-week-old male Sprague-Dawley rats were used in the experiment. AT1R-ECII (0.4 μg/g) was injected subcutaneously into the back neck every two weeks to complete active immunization, which lasted for three months. Ad-BKα-GFP (1×10^10^ pfu/kg) and antagomir-339-3p (20 nmol/rat) injection started in the 6^th^ week of active immunization, and intravenous injection into the rat tail was administered every two weeks until the end of active immunization. All animals used in the experiment were approved by the Animal Protection Ethics Committee of Capital Medical University (Ethics Number: AEEI-2014-062). Finally, the tissue was removed after intraperitoneal injection of 20% pentobarbital sodium at a dose of 40 mg/kg.

### Functional testing

A BP-98A animal noninvasive sphygmomanometer (Softron, Japan) was used to monitor the blood pressure of awake model rats. Before formally testing and recording the testing data, the model animals were allowed to adapt to the pressure stimulation of the sphygmomanometer tail cuff every day for one week in advance. Then the arterial blood pressure of the model rats can be monitored.

After anaesthetizing the model rats with 3% isoflurane, a Vevo LAB small animal ultrasound system (Visualsonics, USA) was used to detect the vascular wall thickness and lumen diameter of the thoracic aortas of the model rats.

### Primary culture and subculture

VSMCs were cultured in low glucose DMEM containing 1% penicillin streptomycin and 10% foetal bovine serum. Human embryonic kidney 293A (HEK293A) cells were purchased from ATCC (Manassas, VA) and cultured in high glucose DMEM containing 10% foetal bovine serum. All cells were cultured in a 37°C, 5% CO_2_ cell incubator. All types of cells need to be passaged approximately every 48 h, and before various treatments, cells need to be cultured in serum-free medium for 24 h.

### Enzyme-linked immunosorbent assay (ELISA)

The content of the target inflammatory cytokine in the test sample was detected by kits (Invitrogen, 88-50625, 88-6010, 88-7340, USA). The sample to be tested was incubated with a biotin-labelled antibody, avidin-labelled HRP, and substrates A and B. Finally, stop solution was added. The colour depth of the liquid is proportional to the concentration of the substance to be tested in the sample.

The OD value of AT1-AA in the serum of actively immunized rats needs to be tested regularly to confirm the success of active immunization. The bottom of the 96-well plate was coated with AT1R extracellular second loop antigen peptide in advance and placed at 4°C overnight. The subsequent steps are similar to the kit process.

### Western blotting

The cell and tissue protein concentrations were evaluated by a BCA protein detection kit (Thermo, 23227, America). A 10% or 12% SDS-PAGE gel was used to separate the target protein from the total protein (20 μg) and then transferred the target protein to a PVDF membrane. The cells were blocked with 5% skimmed milk at room temperature for 1 h and then incubated with the corresponding primary antibody at 4°C overnight. After washing with TBST the next day, the PVDF membranes were incubated with 1:4000 diluted secondary antibody and developed with ECL reagent after washing with TBST.

### RT-PCR

TRIzol and RNA extraction kits were used to extract total mRNA and microRNA from vascular tissue or cultured VSMCs, and reverse transcription into cDNA was performed using a reverse transcription system (Thermo, K1622, USA for total mRNA and GenePharma, China for microRNA). Next, cDNA was amplified according to the amplification system. Finally, the content of amplified genes in the original sample was calculated and analysed based on the Ct value in the amplification result.

### Transfection of plasmid, miRNA and siRNA

The NEDD4L-5’UTR and BKα-3’UTR plasmids were entrusted to Beijing Likely Biotechnology Co., Ltd., and their sequences were inserted into the psi-check 2 dual luciferase miRNA target expression vector (Suppl. Figure 4a). Gene Pharma designed and synthesized the miR-339-3p mimic, miR-339-3p inhibitor, control irrelevant sequences, and BKα and NEDD4L siRNA sequences. Transfection was performed with Lipofectamine 2000, and after 6 h of transfection, the cells were cultured in serum-free DMEM. The lysed cells were finally collected after the relevant operations were performed according to the experimental requirements, and Western blotting, RT-PCR and dual luciferase activity detection were performed.

### Coimmunoprecipitation

The protein volume required by the experimental system based on the protein concentration was calculated, antibodies and protease inhibitors were added and mixed thoroughly at 4°C for 1 h. Then, agarose beads were added and mixed thoroughly overnight at 4°C. The next day, the agarose beads were washed repeatedly with buffer at 4°C. Finally, after discarding the buffer, the agarose beads were boiled with an equal volume of 2× loading mixture, the protein was separated by SDS-PAGE, and the target protein was detected with the corresponding antibody.

### Protein mass spectrometry

The protein-bound agarose beads obtained by coimmunoprecipitation were sent to the company for protein spectroscopy detection (Suppl. Excel).

### Immunofluorescence

Paraffin sections of the thoracic aorta of model rats were used for immunofluorescence staining. After the sections were deparaffinized and hydrated, the antigen was repaired by the high-pressure method, blocked with 10% goat serum, and then incubated with primary antibodies against BKα (Alomone Labs, APC-107, Israel), BKβ1 (Alomone Labs, APC-036, Israel), CD3 (Abcam, ab5690, UK), CD19 (Bioss, bs-0079R, China), CD68 (Affinity Biosciences., DF7518, USA) and α-SMA (Abcam, ab7817, UK) at 4°C overnight. VSMCs were fixed in 4% paraformaldehyde for 10 min at room temperature. After washing with PBS, the smooth muscle cells were also blocked with 10% goat serum. Finally, the cells were incubated in NEDD4L and α-SMA primary antibodies at 4°C overnight. The next day, the paraffin sections and cells were washed with PBS and incubated with a fluorescein-conjugated secondary antibody for 1 h at 37°C. After thorough washing, the tablets were sealed with a sealing solution containing DAPI. The image was acquired with a confocal microscope.

### Fluorescence in situ hybridization (FISH)

For miR-339-3p fluorescence in situ hybridization (FISH), cells were fixed in 4% paraformaldehyde for 15 min at room temperature and washed with PBS after infiltration with 0.1% Triton X-100. The process was carried out using a Gene Pharma fluorescence in situ hybridization kit. Then, according to the operation procedure of the kit, the miR-339-3p red fluorescent (5’ end and 3’ end were both labelled with CY3) probe was used to detect the expression of miR-339-3p in VSMCs.

### Dual-luciferase reporter assay

The test was performed according to the operating instructions of the dual luciferase activity test kit (Vazyme Biotech, DL101-01, USA). The reagents in the kit were diluted and prepared in advance. After washing the cells with PBS, 100 μl diluted lysis buffer was added to each well, and the culture plate was shaken with a shaker for 15 min at room temperature. The lysate was centrifuged, 20 μl supernatant was added to each well of the test plate, 100 μl Luciferase Assay Buffer II (LAR II) was added to detect firefly luciferase activity, and 100 μl stop solution was added to detect Renilla luciferase activity. The ratio of the fluorescence intensity of fireflies to the fluorescence intensity of Renilla reflects the relative fluorescence value of each group.

### Selection of AT1-AA monoclonal antibody cell line

The hybridoma cells were made from the human AT1R extracellular second loop sequence. The hybridoma cells in good condition and able to bind to the extracellular second loop of AT1R were injected into the abdomen of mice to produce ascites. After AT1-AA extraction, the purity and activity of the resulting ascites were detected, including the detection of antibody light and heavy chain (Suppl. Figure 1D), increased beating of neonatal rat cardiomyocytes (Suppl. Figure 1E) and vascular ring detection of vasoconstriction (Suppl. Figure 1F). Finally, select the cell line that allows mice to produce active AT1-AA for follow-up research.

### Preparation of monoclonal AT1-AA

The selected hybridoma cells are cultured, and monoclonal AT1-AA is obtained from the culture supernatant. Hybridoma cells were grown in 1640 RPMI medium containing 8% fetal bovine serum. Select hybridoma cells with good growth status and suitable growth rate, and collect the culture supernatant by centrifugation. The culture supernatant of hybridoma cells was filtered with 0.45 μm filters, and the IgG in the culture supernatant of hybridoma cells was purified with a protein G affinity chromatography column, which is the AT1-AA required for our experiment.

### Isolation and identification of circulating extracellular vesicles in rats

Ultracentrifugation is the most commonly used method for purification of extracellular vesicles. Use low-speed centrifugation and high-speed centrifugation to alternately separate vesicles of similar size. The whole process includes centrifugation at 300 g for 10 min, centrifugation at 2000 g for 10 min, and centrifugation at 10,000 g for 30 min. So far, the supernatant has been retained at each step. Finally, the supernatant was collected, centrifuged at 100,000 g for 70 min, and centrifuged twice to collect the precipitate. The precipitate was an extracellular vesicle. The extracted extracellular vesicles were observed under electron microscope, the particle diameter was detected by Nanoparticle Tracking Analysis (NTA), and the surface markers CD9, CD81 and calnexin of the extracellular vesicles were detected to identify the isolated particles as extracellular vesicles.

### Statistical Analysis

We used GraphPad Prism 8 and SPSS 26 software to draw and analyse the data. All the data are presented as the mean ± standard error (SEM). The differences in normally distributed data were analysed using independent sample *t*-tests (two groups) or one-way ANOVA (> 2 groups). Pearson test is used for correlation analysis. A value of p < 0.05 was considered statistically significant.

## Supplementary Material

Supplementary figures are visible in additional documentation.

### Supplementary Information

#### Supplementary figures and figure legends

**Suppl. Figure 1.**
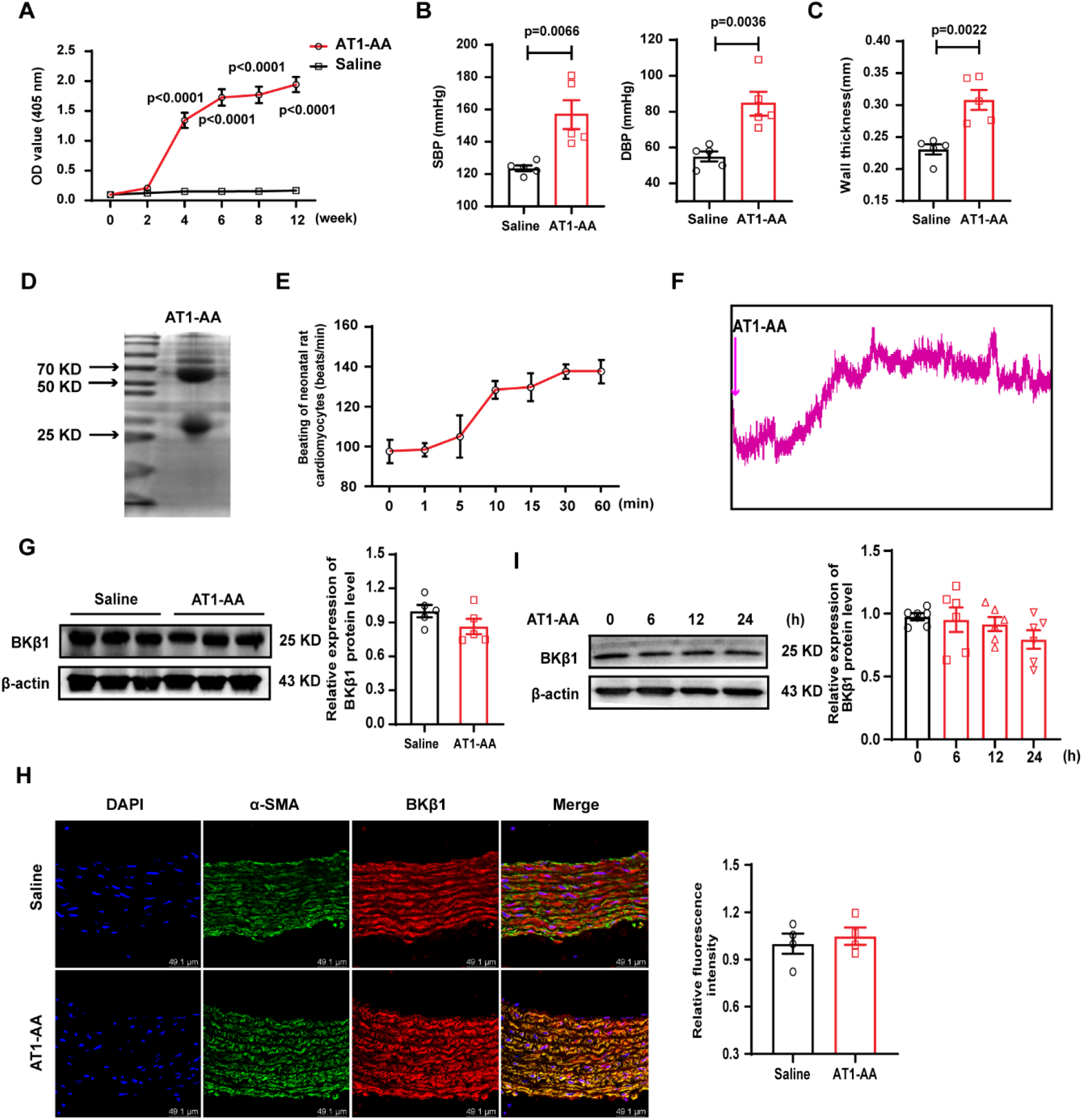
Successful establishment of AT1-AA-positive model does not affect the expression of BKβ1 protein. (A) 12 weeks after active immunization with AT1R-ECII, the OD value of serum AT1-AA in rats was detected. Detection of (B) caudal vein blood pressure and (C) vascular wall thickness of thoracic aorta in AT1-AA-positive rats, n=5. (D) The light and heavy chains of AT1-AA isolated by SDS-PAGE gel. (E) Effect of AT1-AA on the beating of neonatal rat cardiomyocytes. (F) The effect of AT1-AA on vasomotion was detected by vascular ring. Detection of BKβ1 protein level in (G) thoracic aorta of AT1-AA-positive rats and (I) VSMCs treated with AT1-AA, n=5. (H) Detection of BKβ1 protein level in thoracic aorta of AT1-AA-positive rats by immunofluorescence method, n=4.

**Suppl. Figure 2.**
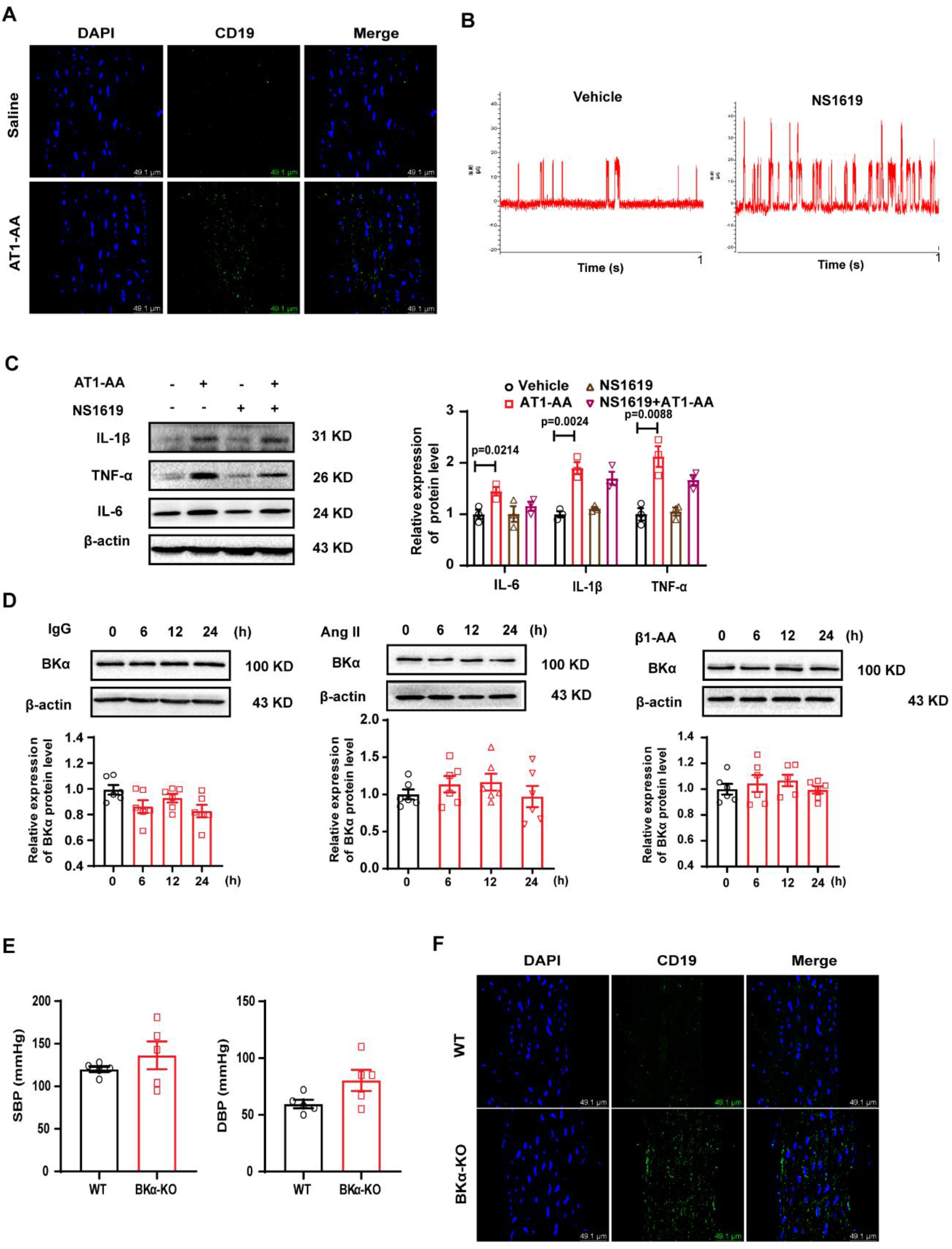
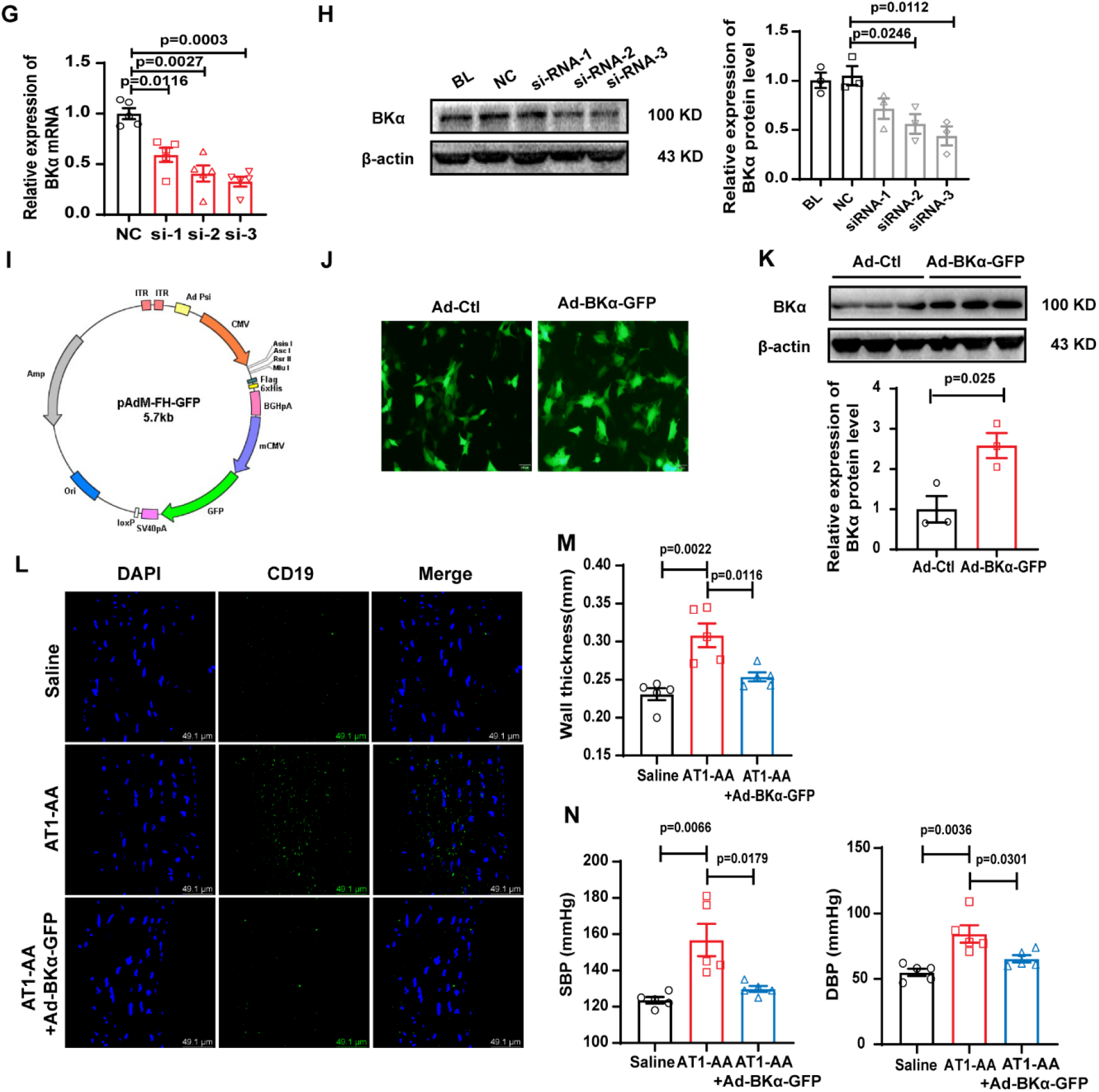
The function and expression of BKα are involved in the inflammatory phenotype of VSMCs aggravated by AT1-AA. The infiltration of inflammatory cells in the vascular wall of the thoracic aorta of AT1-AA-positive rats was detected by (A) immunofluorescence, bar=49.1 μm. (B) The open frequency of BK channel increased after SMCs were treated with NS1619. (C) The expression of inflammatory cytokines was detected after VSMCs were treated with NS1619, n=3. (D) The expression of inflammatory cytokines was detected after VSMCs were treated with IgG, β1-AA and Ang II, n=6. (E) Detection of systolic and diastolic blood pressure in BKα-KO rats. (F) Immunofluorescence was used to detect the infiltration of inflammatory cells in the blood vessel wall of the thoracic aorta of BKα-KO rats. Screening of effective siRNA sequences of BKα through (G) mRNA and (H) protein level, n=3. (I) BKα virus vector map. (J) Photographing the infection efficiency of primary thoracic aortic VSMCs in rats 24 h after BKα overexpression adenovirus infection. (K)Detection of BKα protein expression in SD rats infected by BKα adenovirus, n=3. (L) Immunofluorescence were used to detect the infiltration of inflammatory cells in the blood vessel wall of BKα-overexpression rats. (M) Thoracic aorta wall thickness and (N) tail blood pressure of model rats were measured after BKα adenovirus infection in AT1-AA-positive rats, n=5.

**Suppl. Figure 3.**
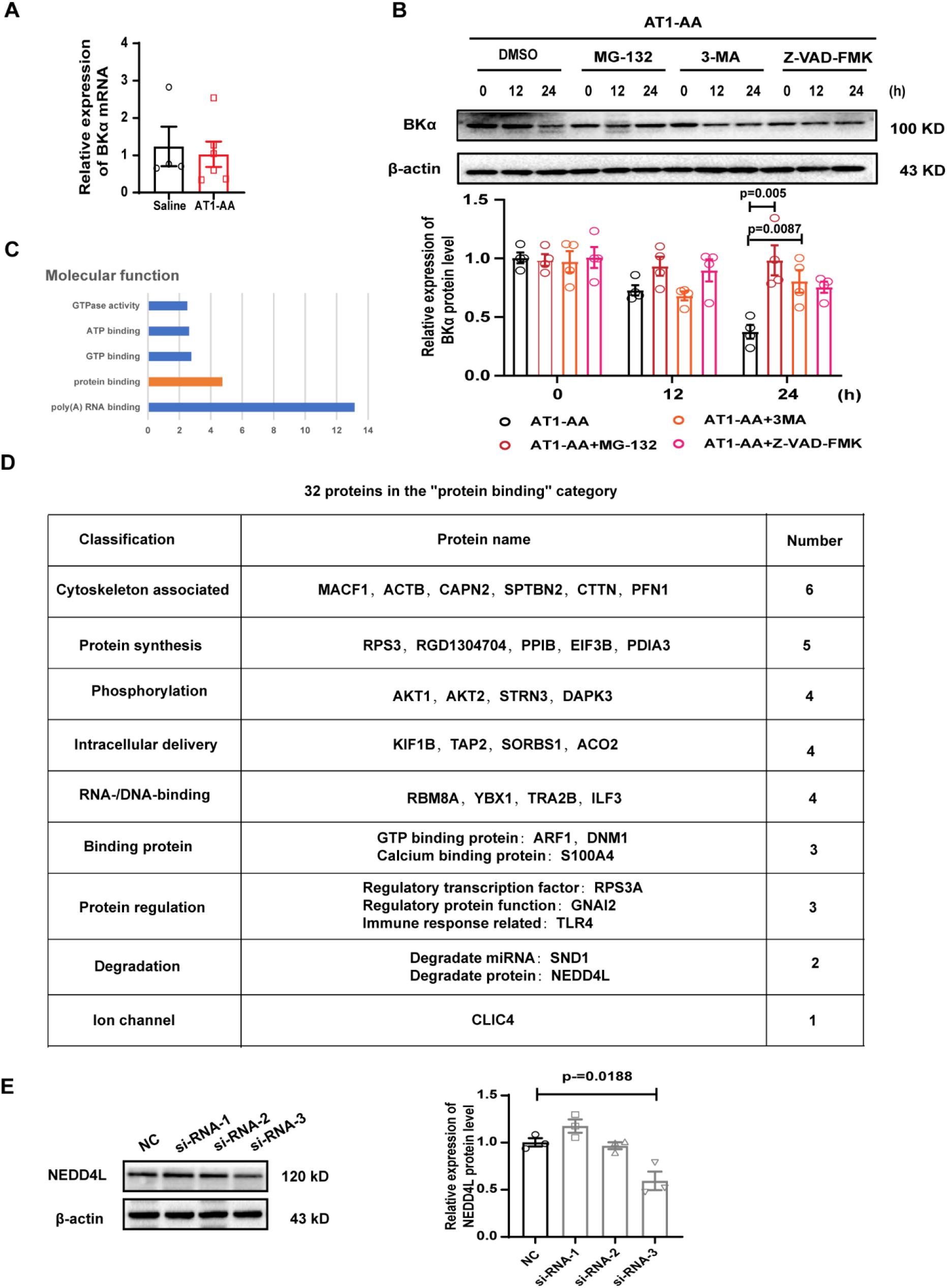
Screening NEDD4L by protein profile and selecting effective NEDD4LsiRNA screening. (A) Detection of BKα mRNA expression in the aorta of AT1-AA positive rats by PCR. (B) The possible pathway which AT1-AA downregulates the BKα protein level in VSMCs was detected by Western blot, n=4. (C) Use protein profile to screen NEDD4L, a protein related to ubiquitination in the protein binding category. (D) Use uniprot database to analyze the functions of 32 proteins in the “protein binding” category. (E) NEDD4L protein level was analyzed by western blot. Data are presented as mean ± SEM, n=3. Statistical analyses by independent-samples t test and one-way ANOVA.

**Suppl. Figure 4.**
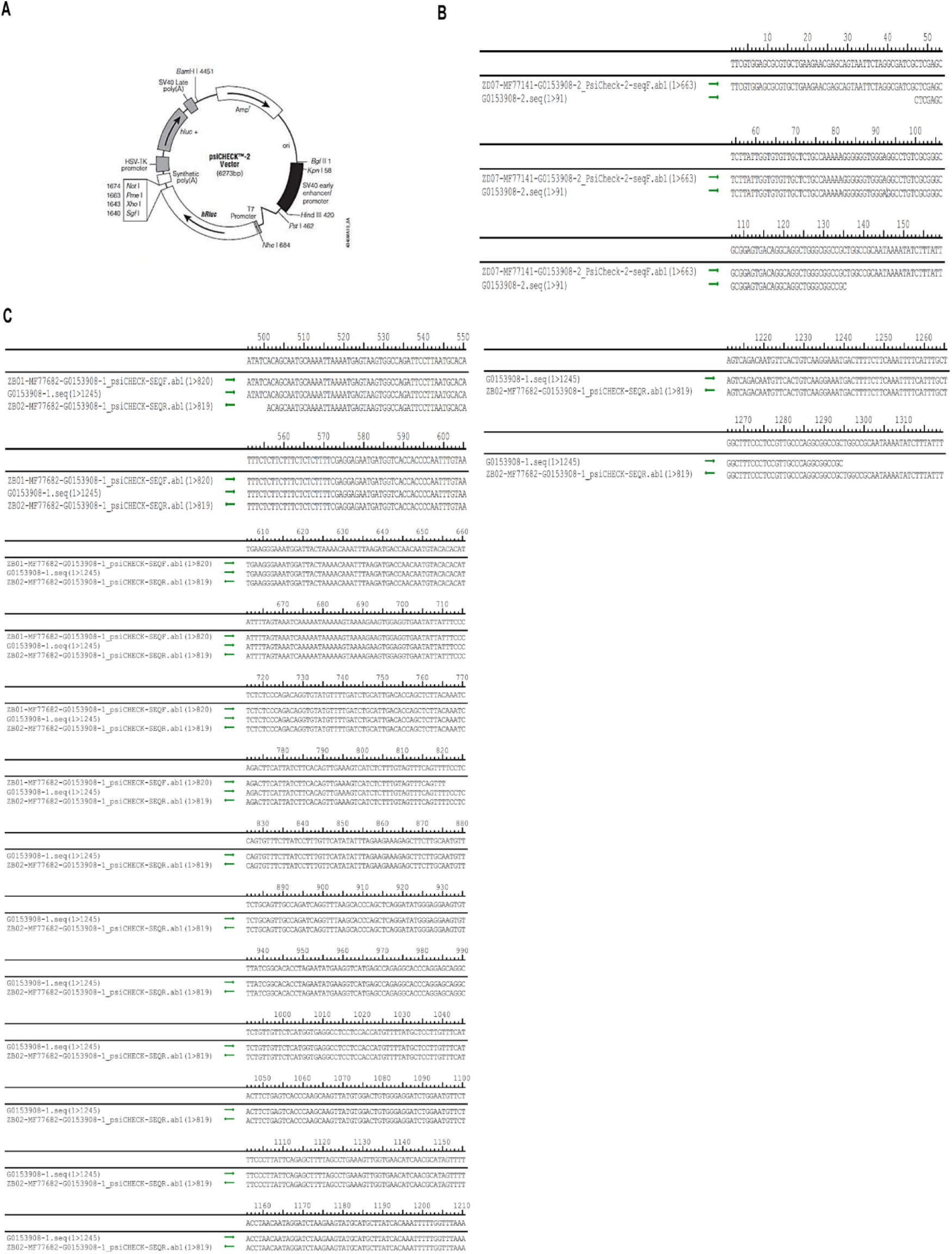
Plasmid vector map and plasmid sequence. (A) Psi-check 2 plasmid vector map. (B) NEDD4L 5’UTR and (C) BKα 3’UTR plasmid sequence.

**Suppl. Figure 5.**
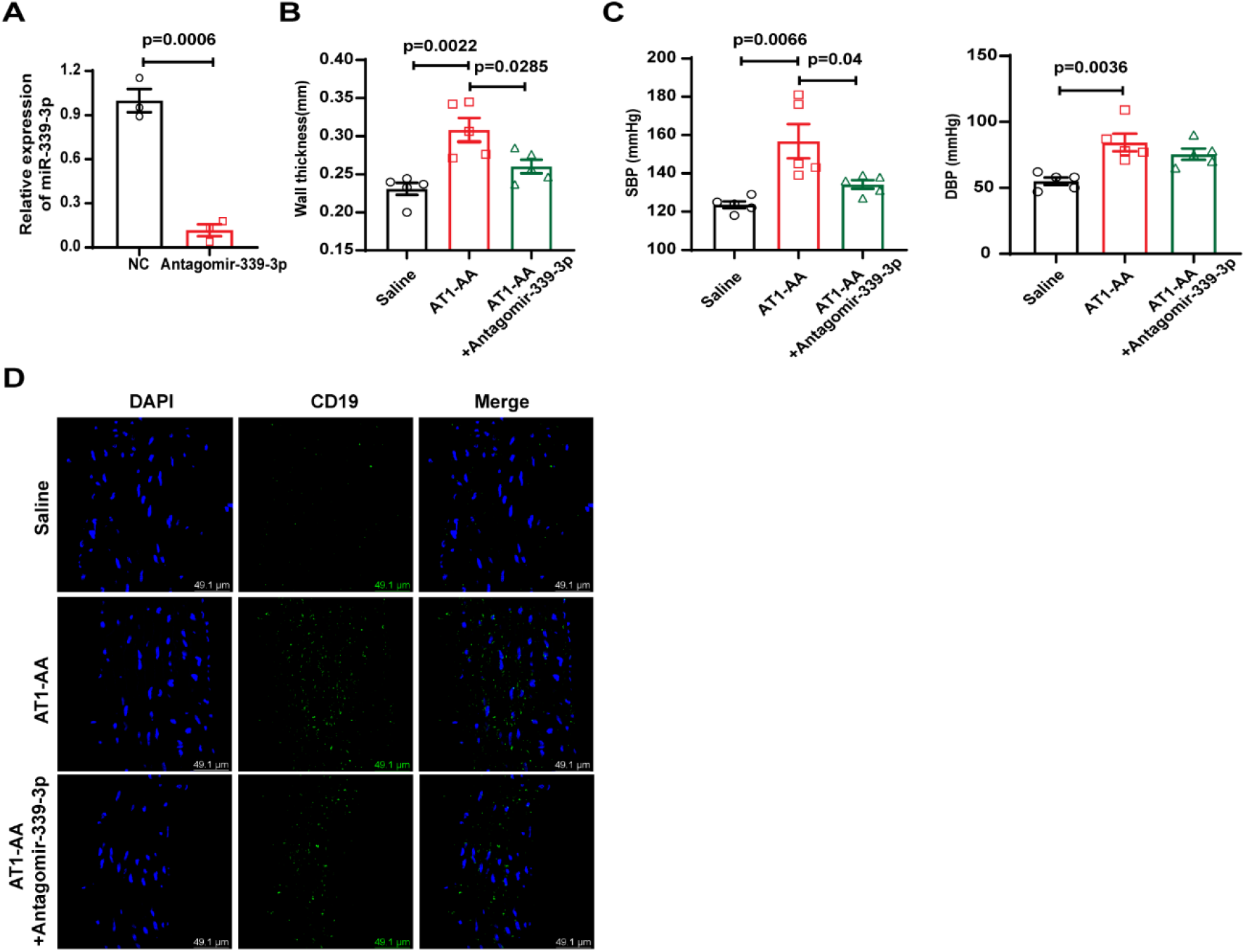
Antagomir-339-3p reversed the vascular injury in AT1-AA-positive rats. (A) Antagomir-339-3p significantly down-regulated the content of miR-339-3p in VSMCs, n=3. (B) Thoracic aorta wall thickness and (C) tail blood pressure of model rats were measured after antagomir-339-3p infection in AT1-AA-positive rats, n=5. (D) The marker of B lymphocytes surface was detected via immunofluorescence method, bar=49 μm.

